# ASPYRE-Lung: Validation of a simple, fast, robust and novel method for multi-variant genomic analysis of actionable NSCLC variants in tissue

**DOI:** 10.1101/2024.02.07.579135

**Authors:** Ryan T Evans, Elizabeth Gillon-Zhang, Julia N. Brown, Katherine E. Knudsen, Candace King, Amanda S Green, Ana-Luisa Silva, Justyna M. Mordaka, Rebecca N. Palmer, Alessandro Tomassini, Alejandra Collazos, Christina Xyrafaki, Iyelola Turner, Chau Ha Ho, Dilyara Nugent, Jinsy Jose, Simonetta Andreazza, Kristine von Bargen, Eleanor R. Gray, Magdalena Stolarek-Januszkiewicz, Aishling Cooke, Honey Reddi, Barnaby W Balmforth, Robert J Osborne

## Abstract

Genomic variant testing of tumors is a critical gateway for patients to access the full potential of personalized oncology therapeutics. Current methods such as next-generation sequencing are costly and challenging to interpret, while PCR assays are limited in the number of variants they can cover. We developed ASPYRE® (Allele-Specific PYrophosphorolysis REaction) technology to address the urgent need for rapid, accessible and affordable diagnostics informing actionable genomic target variants of a given cancer. The targeted ASPYRE-Lung panel for non-small cell carcinoma covers 114 variants in 11 genes (*ALK, BRAF, EGFR, ERBB2, KRAS, RET, ROS1, MET & NTRK1/2/3*) to robustly inform clinical management. The assay detects single nucleotide variants, insertions, deletions, and gene fusions from tissue-derived DNA and RNA simultaneously. We tested the limit of detection, specificity, analytical accuracy and analytical precision of ASPYRE-Lung using FFPE lung tissue samples from patients with non-small cell lung carcinoma, variant-negative FFPE tissue from healthy donors, and FFPE-based contrived samples with controllable variant allele fractions. The sensitivity of ASPYRE-Lung was determined to be ≤ 3% variant allele fraction for single nucleotide variants and insertions or deletions, 100 copies for fusions, and 200 copies for MET exon 14 skipping. The specificity was 100% with no false positive results. The analytical accuracy test yielded no discordant calls between ASPYRE-Lung and expected results for clinical samples (via orthogonal testing) or contrived samples, and results were replicable across operators, reagent lots, runs, and real-time PCR instruments with a high degree of precision. The technology is simple and fast, requiring only four reagent transfer steps using standard laboratory equipment (PCR and qPCR instruments) with analysis via a cloud-based analysis algorithm. The ASPYRE-Lung assay has the potential to be transformative in facilitating access to rapid, actionable molecular profiling of tissue for patients with non-small cell carcinoma.

## Introduction

Worldwide, over 2 million people are diagnosed with lung cancer, which has the highest mortality rate of any cancer (1). In particular, non-small cell lung carcinoma (NSCLC) has a five-year survival rate of just 16% when patients with metastatic NSCLC are treated with chemotherapy alone (2). Historically, standard of care treatment for NSCLC included platinum-based cytotoxic chemotherapy. Prognosis has significantly improved following the emergence of targeted therapies, which typically inactivate oncogenic growth factors and their receptors or inhibit oncogenic tyrosine kinase pathways (3). In addition to higher therapeutic success rates, targeted therapies are often better tolerated, with reduced side effects in patients thus improving quality of life (4). There are now over 30 FDA-approved targeted therapies for NSCLC (5), each targeting specific drivers of this disease, making treatment personalized to the genomic variants of a patient’s tumor.

In order to identify patients who are most likely to benefit from specific targeted therapeutics, tools that enable the detection of multiple variants from a single small quantity patient sample are required. Small core needle-biopsies yielding limited material are becoming increasingly common, and repeated invasive specimen collection from patients for comprehensive genomic testing is not feasible for many reasons (including safety and access to tissue). PCR and fluorescence in-situ hybridization are commonly utilized diagnostic tools. However, both methods test only a limited number of mutations, quickly leading to sample exhaustion, and thus limiting the opportunity to identify the most appropriate targeted therapeutic option. Conversely, next-generation sequencing (NGS) can analyze multiple variations from a single sample but is a costly diagnostic tool, with complex laboratory processes, a suboptimal turnaround time (TAT), and the need for comprehensive bioinformatics analysis, which can be difficult to interpret even in a comprehensive cancer care setting. Moreover, as only a small subset of the mutations detected by NGS are clinically actionable, detection of such a broad panel (including variants of unknown significance) lends little benefit and added complexity to clinical decision making. While patients wait for diagnostic results from molecular testing, non-targeted standard cytotoxic chemotherapies are frequently administered to mitigate tumor progression and address patient safety, which can compromise outcomes even if a prelude to targeted therapy (4). Similarly, guidelines indicate that appropriate targeted therapy should take precedence over treatment with an immune checkpoint inhibitor as concurrent or sequential immunotherapy and targeted therapy can lead to toxicity (6). Overall, fewer than 50% of patients with NSCLC receive appropriate therapy due to either not being tested, not receiving molecular variant testing results in a timely manner, or not receiving appropriate treatment (7). Life-saving targeted therapies thus remain underutilized. Taken together, it is clear that access to rapid, robust actionable genomic testing is critical in enabling patients to access the appropriate treatment in a timely manner.

To address this genomic testing gap, we have recently described ASPYRE, a new technology for the detection of DNA variants and RNA fusions (8,9). Previously, we have shown that ASPYRE technology is highly sensitive, allowing the detection of specific DNA sequences to single-molecule level (8) and RNA fusion detection of under six copies (9), alongside high specificity and robustness against interference from carryover contaminants from formalin fixed paraffin embedded (FFPE) samples (8,9).

The ASPYRE assay comprises four sequential enzymatic stages with reagent transfer between each stage (Figure 1). Briefly, mutant and wild-type molecules are amplified by PCR (or RT-PCR), and the remaining DNA polymerase is enzymatically digested. Amplicons are made single-stranded via exonuclease digestion. Probes that perfectly match target variants hybridize to mutant targets and are subject to pyrophosphorolysis using a DNA polymerase with no exonuclease activity. Conversely, probes that hybridize to wild-type targets are mismatched, and the mismatch prevents pyrophosphorolysis past the variant site. Probes that have been subject to pyrophosphorolysis beyond the variant site are circularized by ligation and amplified by an isothermal hyper-branched rolling-circle amplification reaction that is multiplexed to detect amplification in four fluorescence channels. The ASPYRE assay is run on thermal cyclers and quantitative real-time PCR (qPCR) instruments, which are readily available in most clinical laboratories.

**Figure 1.**
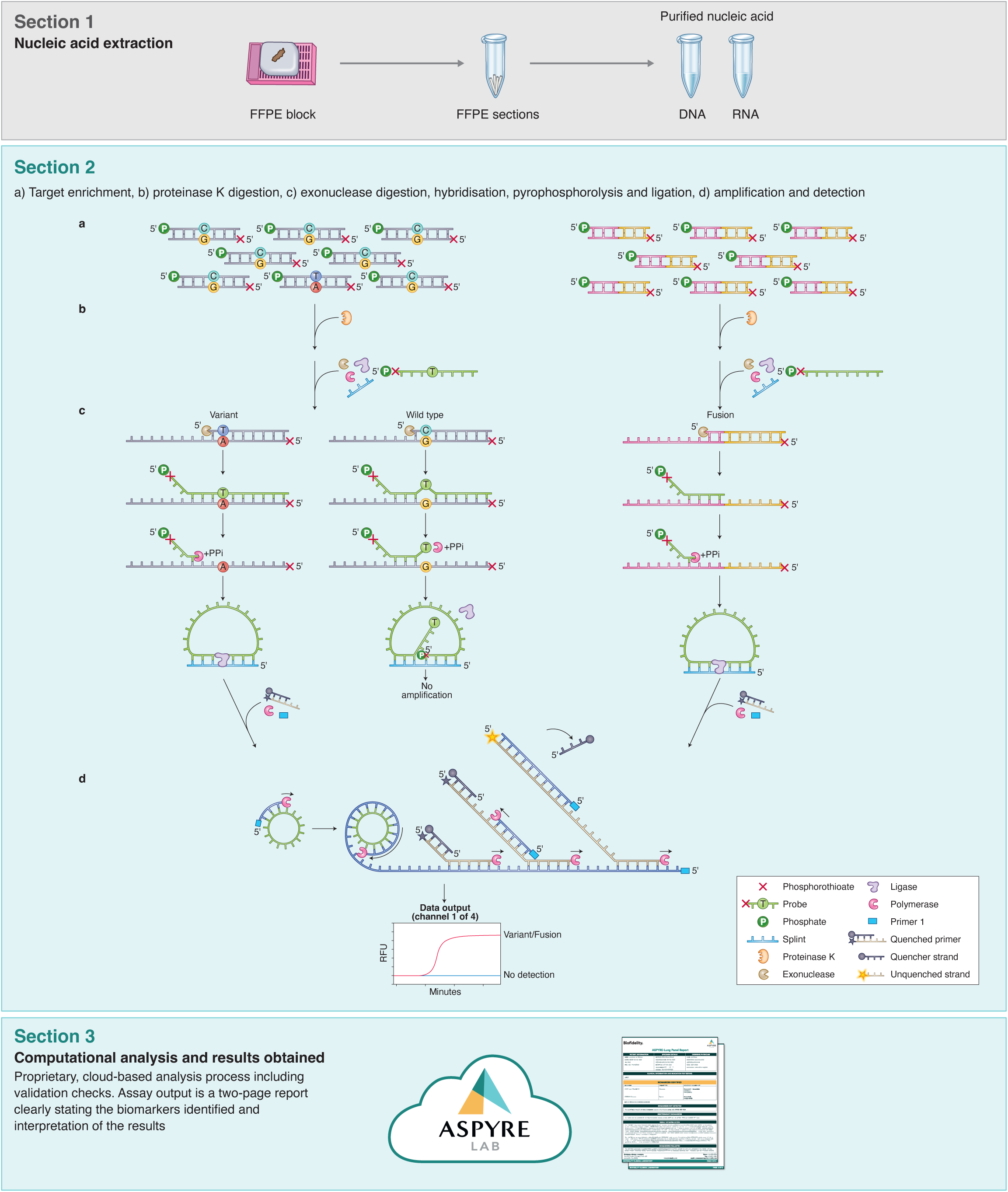
Parallelised workflow of the ASPYRE-Lung targeted mutation panel. Workflow schematic, showing the four separate steps of the assay for DNA (Section 2, left) and RNA (Section 2, right). While steps shown in section 2 differ technically between DNA and RNA, there is no difference to the user in handling the assay, facilitating ease of use.

Herein, we describe the ASPYRE-Lung assay and its performance in our Clinical Laboratory Improvement Amendments (CLIA)-certified laboratory. ASPYRE-Lung enables the concurrent detection of 77 DNA variants, 36 RNA fusions and *MET* exon 14 skipping (S1 Table). Importantly, the variants detected within the assay are clinically actionable and recommended by the National Comprehensive Cancer Network (NCCN), College of American Pathologists (CAP), Association for Molecular Pathology (AMP) and European Society of Medical Oncology (ESMO) advanced NSCLC treatment guidelines. The assay uses DNA and RNA extracted from FFPE tissue samples. The TAT from sample extraction to analysis can be completed within two days. A single patient sample is analyzed in a total of 24 wells, allowing 16 patient samples to be run on a 384-well instrument.

We assessed the performance of the ASPYRE-Lung assay across a range of parameters using both contrived samples and FFPE patient lung tissue. The assay limit of detection (LoD) was determined using serially diluted contrived DNA and RNA samples, and assay specificity (Limit of Blank, LoB) was established using FFPE variant-negative tonsil tissue samples from patients without a known cancer diagnosis. Analytical accuracy was assessed by determining variant calls from FFPE NSCLC lung tumor tissue samples, and analytical precision assessed using FFPE NSCLC lung tissue and variant-negative FFPE tissue. Additionally, common exogenous contaminants (potential interfering substances) from nucleic acid extraction were added to DNA and RNA samples to determine the effect on assay performance. Overall, this study successfully validated the performance of the ASPYRE-Lung assay on FFPE lung tissue samples in our CLIA-certified laboratory. Importantly, ASPYRE-Lung will address the gap clinicians face in obtaining sensitive and actionable mutation testing for patients with NSCLC, at a lower cost, faster TAT, and with minimal sample requirements.

## Results

### The ASPYRE-Lung assay analyzes DNA and RNA derived from tissue

The ASPYRE-Lung assay assesses the status of 114 actionable variants across 11 genes from paired DNA (20 ng) and RNA (6 ng) derived from FFPE lung tissue in a single workflow. Multiplexed targeted PCR amplification of 9 exons of DNA (KRAS exons 2 & 3, BRAF exon 15, ERBB2 exons 17 & 20, EGFR 18, 19, 20, and 21) occurs alongside a separate but parallel RT-PCR reaction to amplify any of 36 RNA fusion targets and one exon skipping event that are present in the sample. The digestion, hybridisation, pyrophosphorolysis and isothermal amplification steps are common to both DNA and RNA. The result for each variant detected by the ASPYRE-Lung panel is interpreted from one of four fluorescent channel signals within each of the 20 DNA and 2 RNA wells (these also incorporate a positive control well for DNA and RNA respectively). An additional two wells are used as negative controls for the DNA and RNA detection reactions. The fluorescent output from the qPCR instrument is analyzed through ASPYRELab software to provide variant call results for each sample, including assay and sample quality checks. Combined with patient information, the parsed results are used to generate a clinical report that indicates presence or absence of a specific mutation (e.g. BRAF exon 15 p.V600E) or class of mutation (e.g. ALK fusion) and the potential therapeutic options available, without manual bioinformatic interpretation. An assay schematic is illustrated in Figure 1.

**The limit of detection of ASPYRE-Lung** is ≤ 3% VAF for SNV and indels, ≤ 100 copies for fusions, and ≤ 200 copies for MET exon 14 skipping

The LoD95 was established as the lowest test level with at least a 95% hit rate. This was determined in two stages: first by estimating the value within a wide range of VAFs or copies for each class-representative variant in the panel (DNA single nucleotide variations (SNV), DNA insertions or deletions (indel), RNA fusion, and *MET* exon 14 skipping), and second by confirming the estimated value with greater replicate testing power. During the estimation stage all replicates were positive across all SNV and indel samples at 3% VAF, all fusion samples at 100 copies, and the *MET* exon 14 skipping mutation at 200 copies, although one false negative was observed for a single exon 14 skipping replicate at 400 copies (S2 Table). The levels chosen for the estimation phase may therefore have been conservative, and the LoD95 likely lies below, or considerably below, these levels.

Secondly, the LoD95 was confirmed by testing 20 replicates of each variant at the estimated LoD95 using a new reagent lot. A minimum of 17/20 (85%) positive results was required for confirmation per variant as per CLSI EP17-A2 specifications with 20 replicates (an upper one-sided 95% confidence limit of 93.8%). At this confirmation stage, 100% of replicates were positive across all SNV and indel samples at 3% VAF, 100% of *RET*, *ROS1* and *NTRK1/2/3* fusions were positive at 100 copies, and 100% of the *MET* exon 14 skipping mutation at 200 copies (Table 1). 90% (18/20) of replicates of the *ALK* fusion were positive at 100 copies. Therefore, the confirmed LoD95 for SNV and indels was ≤ 3%, for fusions ≤ 100 copies, and for *MET* exon 14 skipping mutation ≤ 200 copies (Table 1). The LoD of each variant was confirmed at the lowest tested level, which suggests that the true LoD95 likely lies below (and possibly considerably below) 3% VAF for SNVs and indels, 100 copies for fusions, and 200 copies for *MET* exon 14 skipping.

**Table 1:** LoD95 confirmation data.

|  |  |  |  |  | Confirmation positive hit rate |  |
| --- | --- | --- | --- | --- | --- | --- |
| Variant type | Gene | Exon | Protein variant | COSMIC ID | Sample (n=20, %) | Aggregated positive/total tests* |
| DNA |  |  |  |  | 3% VAF |  |
| SNV | KRAS | 2 | G12C | COSM516 | 100 | 80/80 |
|  | EGFR | 21 | L858R | COSM6224 | 100 |  |
|  | EGFR | 20 | T790M | COSM6240 | 100 |  |
|  | BRAF | 15 | V600E | COSM476 | 100 |  |
| Deletion | EGFR | 19 | E746_<br>A750del | COSM6223 | 100 | 60/60 |
| Insertion | ERBB2 | 20 | Y772_ | COSM20959 | 100 |  |
|  |  |  | A775dup |  |  |  |
|  | EGFR | 20 | A767_<br>V769dup | COSM12376 | 100 |  |
| RNA Fusions |  |  |  |  | 100 copies |  |
| Fusion | EML4-ALK | E13_A20 | NA | COSF408 | 90 | 118/120 |
|  | KIF5B-RET | K15_R12 |  | COSF1232 | 100 |  |
|  | CD74-ROS1 | C6_R34 |  | COSF1200 | 100 |  |
|  | TMP3-NTRK1 | T8_N10 |  | COSF1329 | 100 |  |
|  | QKI-NTRK2 | Q6_N16 |  | COSF1446 | 100 |  |
|  | ETV6-NTRK3 | E5_N15 |  | COSF571 | 100 |  |
| RNA Exon Skipping |  |  |  |  | 200 copies |  |
| Exon skipping | MET | 14 | L982_D1028<br>del | COSM29312 | 100 | 20/20 |

Results of testing 20 replicate contrived samples at levels determined by the LoD estimation (S2 Table). Confirmed LoD95 by mutation class (SNV, indel, fusion and MET exon 14 skipping) were $\leq 3\%$ VAF for SNV and indel, $\leq 100$ copies for fusions, and $\leq 200$ copies for MET exon 14 skipping. \*Results were aggregated across the given variant class. NA, not applicable.

### The ASPYRE-Lung assay in FFPE tissue is highly specific

Next, the specificity of the assay was tested using DNA and RNA extracts from 30 FFPE tonsil samples from donors with no history of cancer diagnosis. DNA and RNA from each FFPE sample was extracted, and each nucleic acid tested in duplicate using two different reagent lots for a total of 60 tests. One DNA sample was repeated once to obtain a valid result as the internal positive control failed in the first run. There were no positive calls for any sample at any of the 6840 variants analyzed by ASPYRE-Lung (Table 2); therefore, the false positive rate was 0% (0-6% Clopper-Pearson 95%CI) and the LoB of the test is zero (0-4.1% Jeffreys 95% confidence interval).

**Table 2:** Variants tested per sample during the LoB assessment.

| Category | Number (n) |  |  |
| --- | --- | --- | --- |
|  | Tested per ASPYRE assay | Total Tested | Positive |
| Samples | 1 | 60 | 0 |
| Total nucleotide variants tested | 114 | 6840 | 0 |
| SNVs | 26 | 1560 | 0 |
| Indels + complex substitutions | 31 + 20 | 3060 | 0 |
| Fusions | 36 | 2160 | 0 |
| Exon skipping | 1 | 60 | 0 |
| Reportable variants | 71 | 4260 | 0 |

A total of 60 independently extracted nucleic acid samples from 30 FFPE variant-free blocks were tested for all variants covered by the ASPYRE-Lung panel. A single 'Reportable Variant' may cover multiple nucleotide variants where the associated therapeutics are identical. For example, all gene fusions that involve *NTRK1*, *NTRK2* or *NTRK3* will be reported under a single

### ASPYRE-Lung has high analytical accuracy

The analytical accuracy of the ASPYRE-Lung assay was assessed using both contrived specimens (Table 3 and S3 Table) and nucleic acid extracted from 30 FFPE samples from patients with a diagnosis of NSCLC for which targeted enrichment NGS data were available (Table 3 and S4 Table). Each of the contrived samples was tested once at twice the LoD95. Variants were aggregated according to class, and all results were according to expected output with 100% PPA and 100% NPA across all samples (Table 3).

**Table 3:**
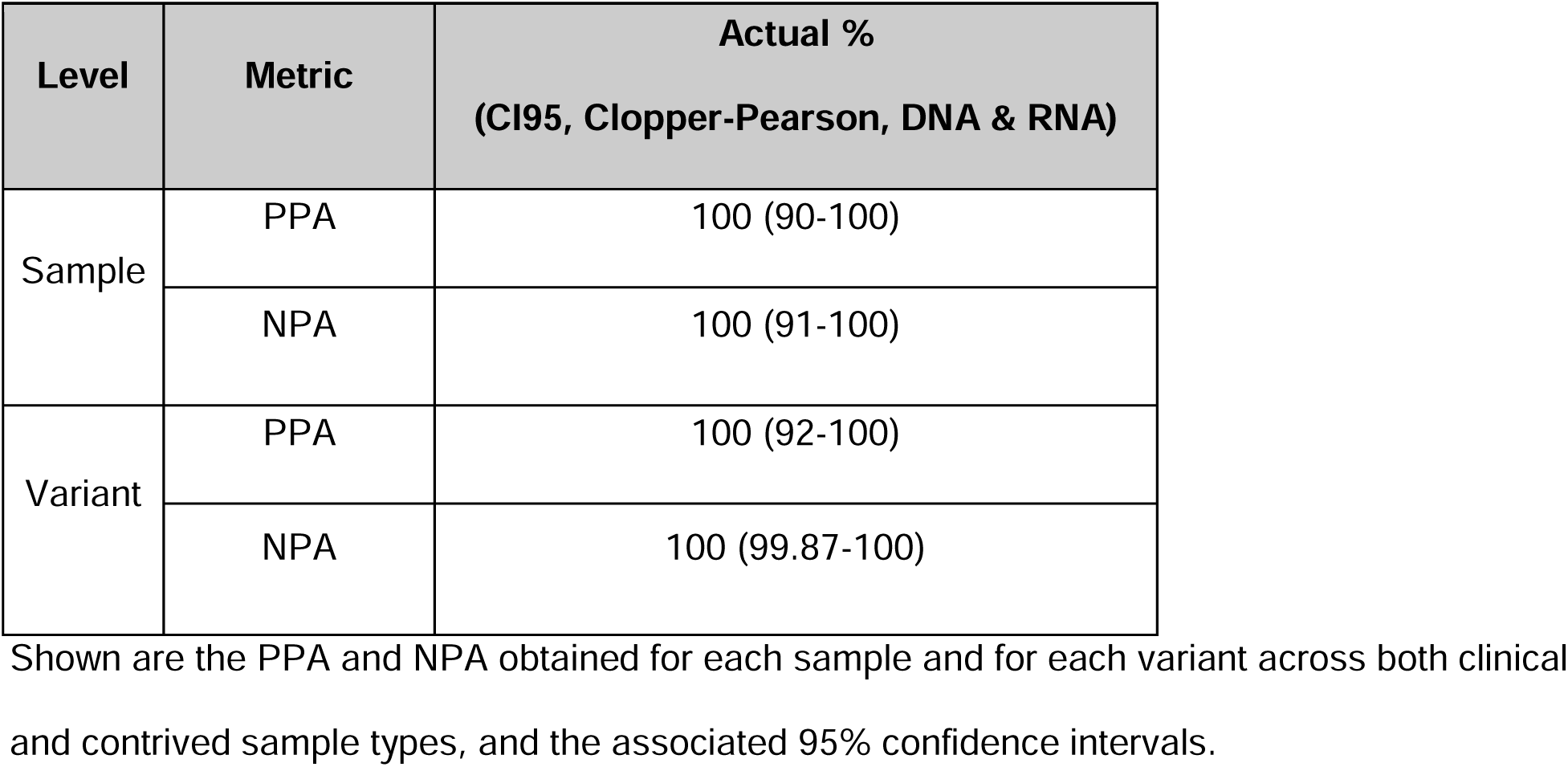
Summary of analytical accuracy of ASPYRE-Lung assessed using contrived and clinical samples.

### The ASPYRE-Lung assay has excellent analytical precision

The reproducibility and repeatability of the assay were tested using FFPE tissue samples (DNA and RNA from variant-positive and variant-negative patient samples) tested in duplicate across four runs with two instruments by two operators over four days (Table 4). The three FFPE NSCLC variant-positive tissue samples were sequenced by targeted enrichment NGS (S4 Table). All replicate runs were matched with each other and expected results, giving 100% PPA and NPA, and 100% reproducibility and repeatability.

**Table 4:** Summary of analytical precision (repeatability and reproducibility) data.

| Level | Metric | Actual %<br>(CI95, Clopper-Pearson) |
| --- | --- | --- |
| Sample | PPA | 100 (86-100) |
|  | NPA | 100 (93-100) |

Shown are the positive and negative percent agreement values between runs of ASPYRE-Lung, demonstrating 100% reproducibility (inter-run precision) and repeatability (intra-run precision). Samples were assayed in four independent runs across four days by two operators using two real-time PCR instruments and two reagent lots. An expanded dataset from which this table is derived is shown in S5 Table.

### ASPYRE-Lung is not affected by common interfering substances present in nucleic acid extracts

As ASPYRE-Lung utilizes isolated nucleic acids as sample input, it is potentially susceptible to interfering substances carried over during DNA and RNA extraction. Common substances used in nucleic acid extraction that often impact molecular assays include guanidinium salts and ethanol. We tested whether the performance of the ASPYRE-Lung assay was affected by the presence of carry-over chemicals by spiking two DNA and two RNA samples with guanidinium thiocyanate and ethanol, with five replicates per sample and spike-in condition (Table 5). The ethanol test level was the same as used by the Foundation Medicine FoundationOne CDx FDA submission (Summary of Safety and Effectiveness Data, P170019), while guanidine thiocyanate levels were chosen from in-house preliminary testing of levels found in samples with incomplete removal of contaminants. None of the contaminants tested showed any effect on assay results at the levels tested and are therefore considered non-interfering (Table 5).

**Table 5:** Contaminants carried over from FFPE sample extraction do not interfere with the ASPYRE-Lung assay.

|  |  |  |  |  |  | TP / total Replicates |  |  |  |
| --- | --- | --- | --- | --- | --- | --- | --- | --- | --- |
|  |  |  |  |  |  | 5 %<br>Ethanol | Guanidine thiocyanate |  |  |
| Analyte | Gene | Exon | Protein Variant | COSMIC ID | VAF/<br>Copies |  | 5 mM | 10 mM | 20 mM |
| DNA | KRAS | 2 | G12C | COSM516 | 6% | 5/5 | 5/5 | 5/5 | 5/5 |
| DNA | EGFR | 21 | L858R | COSM6224 | 6% | 5/5 | 5/5 | 5/5 | 5/5 |
| RNA | EML4-ALK | E13_<br>A20 | NA | COSF408 | 200<br>copies | 5/5 | 5/5 | 5/5 | 5/5 |
| RNA | KIF5B-RET | K15_<br>R12 | NA | COSF1232 | 200<br>copies | 5/5 | 5/5 | 5/5 | 5/5 |

Two DNA and two RNA contrived samples at twice the LoD were spiked with ethanol or guanidinium thiocyanate to mimic carryover from substances used during sample extraction. Shown are the positive calls made compared to total runs from the subsequent ASPYRE-Lung assay runs. NA, not applicable.

## Discussion

In an ideal scenario, patients newly diagnosed with NSCLC would receive results from genomic tests that identify all actionable driver mutations within a short timeframe after confirmed diagnosis, in order to access the appropriate targeted therapy as soon as possible (4). While practice guidelines recommend molecular testing at the outset for all actionable biomarkers (10,11), around 90% of patients are tested for at least one actionable biomarker, only just under half are tested for five or more biomarkers, with many beginning chemotherapy before results from biomarker testing are returned (12,13). Stratification of patients according to the mutation in their tumor can be achieved through a variety of methods, including single-target and multiplex PCR which look for specific alterations, and NGS which captures specific genomic regions and analyzes changes therein. There are advantages and disadvantages to each method. Targeted methods of detection are well established, faster, and more affordable, but can require running multiple sequential assays in order to capture all possible actionable variants, with the potential for tissue exhaustion and extended laboratory time. NGS has a long TAT and yields large amounts of data that require expert bioinformatics interpretation and may generate technically unclear results for some targets (for example, the prediction of gene fusions at the RNA level from DNA sequence in introns, (7,12).

The characteristics of the ASPYRE-Lung assay make it eminently suitable for all patients with NSCLC to undergo mutational profiling, as it is simple and rapid to perform using standard laboratory techniques and equipment (8,9). The panel covers the guideline recommended 77 mutations from DNA and 36 gene fusions and MET exon 14 skipping events in RNA. In this study, we have estimated and confirmed an LoD95 of ASPYRE-Lung in FFPE tissue at ≤ 3% VAF for SNV and indels found in DNA, ≤ 100 copies for RNA fusions and ≤ 200 copies for MET exon 14 skipping. The specificity of the assay was 100%, with no false positive results yielded from 60 independent tests of 30 variant-free FFPE tissue samples using two lots of reagents.

The accuracy of the assay compared with targeted enrichment NGS was 100% PPA and NPA, with no false negative calls, including for samples with more than one mutation present. Repeated testing of six samples (three positive and three negative) across different days, operators, reagent lots, instruments and assay runs gave 100% consistent results. Finally, spiking samples with potential interfering substances that are commonly found in extraction kits, and can be incompletely removed during pre-analytical procedures, did not affect whether correct calls were made.

While these results demonstrate excellent assay performance, work is ongoing to expand the remit of the assay by further testing and validating performance characteristics on a wide range of samples to address unmet clinical need. Lung tissue biopsies can take different forms, including core needle biopsies and samples taken for cytology (pleural effusions, bronchial washings) which do not meet tissue requirements for NGS or sequential PCR testing. Previous work has shown that the ASPYRE assay has robust results down to 1 ng extracted RNA from clinical lung FFPE tissue samples (9), and this work will be updated and expanded to include DNA and a lower range of sample input levels. Further goals include validating the assay and analysis algorithm for circulating nucleic acid extracted from plasma. As new targeted therapies become available and guidelines change, the panel can be expanded to include detection of additional biomarkers, allowing more patients to access the appropriate targeted therapy for which they may best respond.

## Conclusions

1. There are now many targeted therapies available but biomarker testing remains a crucial roadblock to accessing these highly efficacious and well-tolerated treatments in a timely manner. This is largely due to the limitations of existing genomic testing technologies.
2. ASPYRE is a new class of diagnostic technology that leverages a unique series of exquisitely sensitive and specific enzymatic reactions to give the benefits of multi-gene testing yet with rapid TAT, simple bioinformatics, and with easily interpretable clinical decision making (only actionable markers are tested).
3. In this study, we demonstrate that ASPYRE-Lung FFPE Tissue assay has excellent analytical sensitivity at ≤ 3% VAF for SNV and indels from DNA, ≤ 100 copies for gene fusions from RNA, and ≤ 200 copies MET exon 14 skipping from RNA. The assay has high specificity with no false positive results out of 6,840 calls made from variant-free samples. The assay is also highly reproducible and repeatable across different operators, reagent lots, runs, days and qPCR instruments.
4. Obtaining sufficient quantity and quality of tissue samples from lung biopsies for robust genomic analysis can be challenging, and future plans for ASPYRE-Lung include validating the assay on small biopsy and cytopathology samples. Finally, we are developing a version of the assay that will analyze liquid (plasma) biopsy samples as input to provide results for individuals where tissue specimens are not available.

## Methods

### Reference samples

DNA and RNA variants were selected to represent the most common variant in each target exon from *BRAF* exon 15; *EGFR* exons 18, 19, 20, 21; *KRAS* exons 2, 3; and *ERBB2* exons 17, 20; and the most common *ALK, RET, ROS1, NTRK1/2/3* fusions alongside *MET* exon 14 exon skipping (selected variants are listed in S2 Table). Contrived control oligonucleotides made from DNA (SNV, indel) or RNA (fusions, *MET* exon 14 skipping) were manufactured (DNA from Eurofins, Wolverhampton, UK, RNA from IDT, Leuven, Belgium), quantified by digital PCR (QIAcuity Digital PCR system, Qiagen) at the Biofidelity R&D facility (Cambridge, UK), and spiked into DNA or RNA extracted from FFPE variant-free tonsil tissue quantified by dPCR (DNA) or Qubit (RNA), serially diluted to the appropriate concentrations, and immediately frozen at −20°C (DNA) or −80°C (RNA).

### Clinical samples

FFPE variant-free tonsil tissue blocks from patients without any known cancer diagnosis were procured from a commercial biobank retrospectively, sample selection and data access July 1 2022 (Reprocell, Maryland, USA). FFPE lung tissue blocks with a confirmed NSCLC diagnosis were obtained from commercial biobanks (Geneticist, Tissue Solutions, Reprocell, BocaBio, Cureline, VitroVivo) between August 2020 and August 2021.

### Nucleic acid extraction

FFPE blocks were manually sectioned with a microtome (Shandon Finesse, ThermoFisher), producing three 12 μM thick curls, at the Biofidelity R&D facility (Cambridge, UK). DNA and RNA were extracted from specimens in parallel using the Quick-DNA/RNA^TM^ FFPE miniprep kit (Zymo Research). Nucleic acid concentration was determined with a Qubit™ 1xDNA or RNA high sensitivity kit (ThermoFisher) and stored at −80°C until further usage.

### ASPYRE reaction

The ASPYRE reaction has been described for DNA detection (8) and RNA detection (9) and was performed as carried out previously with input levels of 20 ng DNA per PCR reaction and 6 ng RNA per RT-PCR reaction.

### Orthogonal testing of clinical samples

DNA extracted from FFPE lung tissue samples was sequenced through an orthogonal method by targeted enrichment (Roche Avenio Targeted Assay) and sequencing (NextSeq 500, Illumina) by Glasgow Polyomics (University of Glasgow, UK), according to the manufacturer’s guidelines. Analysis was performed by the Roche Sequencing Solutions team.

### Interfering substances

Four contrived reference samples of COSM516 at 6% VAF, COSM6224 at 6% VAF, COSF408 at 200 copies or COSF1232 at 200 copies prepared in background DNA or RNA (extracted from FFPE variant-free tissue) was spiked with molecular biology-grade ethanol (Sigma-Aldrich) or guanidinium thiocyanate (Sigma-Aldrich) at concentrations mimicking potential carryover during the extraction process (ethanol at 5%, guanidine thiocyanate at 5, 10 or 20 mM).

### Data analysis

Data were downloaded from QuantStudio 5 RealTime PCR System (Thermofisher) instruments running Design and Analysis 2 software. The Raw Data CSV produced by this software was analyzed using custom ASPYRELab v1.0.0 software. This cloud-based web application takes the Raw Data CSV as input and provides variant calls and control statuses as output. All variant calling was blinded to results from orthogonal analyses. The ASPYRELab results were then further collated and analyzed using standalone Python scripts.

## Supporting information

Evans et al Supplementary Information

## Acknowledgements

We thank Kasia Anton, Maksym Artomenko, Ethan A. Clark, Seb Dangerfield, Timon Heide, Donald McAusland, Nicola D. Potts, Paulina Powalowska for assistance in the development of ASPYRE-Lung. We acknowledge the work of the Roche Sequencing Solutions team in Mannheim, Germany in assisting with analyses of NGS data.

## Author Contributions

Methodology: RTE, HR, BWB, RJO

Investigation: RTE, ASG, ALS, JMM, MSJ, AC

Conducted experiments: RTE, EGZ, JNB, KEK, CK, ALS, JMM, AC, CX, IT, CHH, DN, JJ, SA, KVB

Formal analysis: RTE, ASG, RNP, AT, AC

Study Supervision: RJO, BB

Writing - Original Draft Preparation: ERG

Writing - Review & Editing: RJO, RTE, BWB, ASG, MSJ, ALS, JMM, ERG, KEK

## Supporting Information Captions

**S1 Table: List of variants covered by the ASPYRE-Lung panel.** There are 77 variants detected by ASPYRE-Lung using DNA, and 37 detected by analysis of RNA. The assay output is 71 calls, as many calls are identical for multiple targets (e.g. “ALK-positive” for the seven gene fusions involving ALK exon 20 that are detected by ASPYRE-Lung).

**S2 Table. Estimation of the LoD95**. Shown are the analyte type (DNA or RNA) and input level: percent positive samples tested at 3, 5 and 10% VAF for those detected in DNA (SNV and indel); 100, 200, and 300 copies for fusions detected by analysis of RNA; and 200, 400, and 800 copies of RNA for MET exon 14 skipping. *Results were aggregated across the given variant class. NA, not applicable.

**S3 Table. Contrived samples generated for this study and tested in the analytical accuracy assessment.** Oligonucleotides bearing sequences matching the variations were quantified using dPCR, and added to a background nucleic acid pool comprising DNA or RNA derived from FFPE tonsil tissue nucleic acid extracts.

**S4 Table. FFPE lung tissue samples from patients with a confirmed NSCLC diagnosis.** Shown are the clinical characteristics associated with each sample, and how each sample was used during the analytical validation. *Fusion breakpoint not predicted to be detected by ASPYRE-Lung.

**S5 Table. Expanded Results from testing analytical precision.** Results from testing nine clinical samples across four runs using two different instruments, two different reagents lots, by two different operators over four days.

