## Supplementary material for "ASPYRE-Lung: Validation of a simple, fast, robust and novel method for multi-variant genomic analysis of actionable NSCLC variants in tissue": Evans et al Supplementary Information

**Supporting Information**

**S1 Table.**

| **Target Source** | **Gene** | **Exon** | **Variants** |
| --- | --- | --- | --- |
| DNA | EGFR | 18 | G719A, G719S, G719C |
|  |  | 19 | Deletions, insertions |
|  |  | 20 | T790M, S768I, C797S, insertions |
|  |  | 21 | L858R, L861Q |
|  | BRAF | 15 | V600E |
|  | KRAS | 2 | G12A, G12C, G12D, G12F, G12R, G12S, G12V, G13A, G13C, G13D, G13R, G13S |
|  |  | 3 | Q61E, Q61H, Q61K, Q61L, Q61R |
|  | ERBB2 | 17 | V659E |
|  |  | 20 | Insertions |
| RNA | ALK |  | Fusions |
|  | ROS1 |  | Fusions |
|  | RET |  | Fusions |
|  | NTRK1 |  | Fusions |
|  | NTRK2 |  | Fusions |
|  | NTRK3 |  | Fusions |
|  | MET | 14 | Exon 14 skipping |

**List of variants covered by the ASPYRE-Lung panel.** There are 77 variants detected by ASPYRE-Lung using DNA, and 37 detected by analysis of RNA. The assay output is 71 calls, as many calls are identical for multiple targets (e.g. “ALK-positive” for the seven gene fusions involving ALK exon 20 that are detected by ASPYRE-Lung).

**S2 Table**

|  |  |  |  |  |  |  | **Positive Hit Rate** | | | |
| --- | --- | --- | --- | --- | --- | --- | --- | --- | --- | --- |
| **Analyte** | **Variant Type** | **Gene** | **Exon** | **Protein variant** | **COSMIC ID** | **VAF (%) or copies** | **LoD Estimation (%)** | **n** | **Confirmation (n=20, %)** | **Aggregated positive /total tests*** |
| DNA | SNV | KRAS | 2 | G12C | COSM516 | 3 | 100 | 5 | 100 | 80/80 |
|  |  |  |  |  |  | 6 | 100 | 5 | - |  |
|  |  |  |  |  |  | 10 | 100 | 5 | - |  |
|  |  | EGFR | 21 | L858R | COSM6224 | 3 | 100 | 5 | 100 |  |
|  |  |  |  |  |  | 6 | 100 | 5 | - |  |
|  |  |  |  |  |  | 10 | 100 | 5 | - |  |
|  |  | EGFR | 20 | T790M | COSM6240 | 3 | 100 | 5 | 100 |  |
|  |  |  |  |  |  | 6 | 100 | 5 | - |  |
|  |  |  |  |  |  | 10 | 100 | 5 | - |  |
|  |  | BRAF | 15 | V600E | COSM476 | 3 | 100 | 5 | 100 |  |
|  |  |  |  |  |  | 6 | 100 | 5 | - |  |
|  |  |  |  |  |  | 10 | 100 | 5 | - |  |
|  | Deletion | EGFR | 19 | E746_A750del | COSM6223 | 3 | 100 | 5 | 100 | 60/60 |
|  |  |  |  |  |  | 6 | 100 | 5 | - |  |
|  |  |  |  |  |  | 10 | 100 | 5 | - |  |
|  | Insertion | ERBB2 | 20 | Y772_A775dup | COSM20959 | 3 | 100 | 5 | 100 |  |
|  |  |  |  |  |  | 6 | 100 | 5 | - |  |
|  |  |  |  |  |  | 10 | 100 | 5 | - |  |
|  | Insertion | EGFR | 20 | A767_V769dup | COSM12376 | 3 | 100 | 5 | 100 |  |
|  |  |  |  |  |  | 6 | 100 | 5 | - |  |
|  |  |  |  |  |  | 10 | 100 | 5 | - |  |
| RNA | Fusion | EML4-ALK | E13_A20 | NA | COSF408 | 100 | 100 | 5 | 90 | 118/120 |
|  |  |  |  |  |  | 200 | 100 | 5 | - |  |
|  |  |  |  |  |  | 300 | 100 | 5 | - |  |
|  |  | KIF5B-RET | K15_R12 | NA | COSF1232 | 100 | 100 | 4 | 100 |  |
|  |  |  |  |  |  | 200 | 100 | 4 | - |  |
|  |  |  |  |  |  | 300 | 100 | 5 | - |  |
|  |  | CD74-ROS1 | C6_R34 | NA | COSF1200 | 100 | 100 | 4 | 100 |  |
|  |  |  |  |  |  | 200 | 100 | 5 | - |  |
|  |  |  |  |  |  | 300 | 100 | 5 | - |  |
|  |  | TMP3-NTRK1 | T8_N10 | NA | COSF1329 | 100 | 100 | 4 | 100 |  |
|  |  |  |  |  |  | 200 | 100 | 5 | - |  |
|  |  |  |  |  |  | 300 | 100 | 4 | - |  |
|  |  | QKI-NTRK2 | Q6_N16 | NA | COSF1446 | 100 | 100 | 5 | 100 |  |
|  |  |  |  |  |  | 200 | 100 | 5 | - |  |
|  |  |  |  |  |  | 300 | 100 | 5 | - |  |
|  |  | ETV6-NTRK3 | E5_N15 | NA | COSF571 | 100 | 100 | 4 | 100 |  |
|  |  |  |  |  |  | 200 | 100 | 4 | - |  |
|  |  |  |  |  |  | 300 | 100 | 5 | - |  |
|  | Exon skipping | MET | 14 | L982_D1028del | COSM29312 | 200 | 100 | 4 | 100 | 20/20 |
|  |  |  |  |  |  | 400 | 75 | 4 | - |  |
|  |  |  |  |  |  | 800 | 100 | 4 | - |  |

**Estimation of the LoD95**. Shown are the analyte type (DNA or RNA) and input level: percent positive samples tested at 3, 5 and 10% VAF for those detected in DNA (SNV and indel); 100, 200, and 300 copies for fusions detected by analysis of RNA; and 200, 400, and 800 copies of RNA for MET exon 14 skipping. *Results were aggregated across the given variant class. NA, not applicable.

**S3 Table**

| **Variant type** | **Gene** | **Exon** | **Protein variant** | **COSMIC ID** | **% VAF / Copies** | **ASPYRE-Lung** |
| --- | --- | --- | --- | --- | --- | --- |
| SNV | KRAS | 2 | G12C | COSM516 | 6 | Matched expected |
|  | EGFR | 21 | L858R | COSM6224 | 6 | Matched expected |
|  | EGFR | 20 | T790M | COSM6240 | 6 | Matched expected |
|  | BRAF | 15 | V600E | COSM476 | 6 | Matched expected |
| Deletion | EGFR | 19 | E746_  A750del | COSM6223 | 6 | Matched expected |
| Insertion | ERBB2 | 20 | Y772_  A775dup | COSM20959 | 6 | Matched expected |
|  | EGFR | 20 | A767_  V769dup | COSM12376 | 6 | Matched expected |
| Fusion | EML4-ALK | E13_A20 | NA | COSF408 | 200 | Matched expected |
|  | KIF5B-RET | K15_R12 |  | COSF1232 | 200 | Matched expected |
|  | CD74-ROS1 | C6_R34 |  | COSF1200 | 200 | Matched expected |
|  | TMP3-NTRK1 | T8_N10 |  | COSF1329 | 200 | Matched expected |
|  | QKI-NTRK2 | Q6_N16 |  | COSF1446 | 200 | Matched expected |
|  | ETV6-NTRK3 | E5_N15 |  | COSF571 | 200 | Matched expected |
| Exon skipping | MET | 14 | L982_D1028del | COSM29312 | 400 | Matched expected |
| Double Mutant | EGFR | 21 | L858R | COSM6224 | 30 | Matched expected |
|  |  | 20 | T790M | COSM6240 | 15 | Matched expected |
|  | EGFR | 19 | E746_A750del | COSM6223 | 30 | Matched expected |
|  |  | 20 | T790M | COSM6240 | 15 | Matched expected |
|  | EGFR | 19 | E746_A750del | COSM6223 | 30 | Matched expected |
|  |  | 20 | C797S | COSM5945664 | 15 | Matched expected |

**Contrived samples generated for this study and tested in the analytical accuracy assessment.** Oligonucleotides bearing sequences matching the variations were quantified using dPCR, and added to a background nucleic acid pool comprising DNA or RNA derived from FFPE tonsil tissue nucleic acid extracts.

**S4 Table**

| **Sample ID** | **Pathology or Clinical diagnosis** | **Stage** | **Result from Roche Avenio Targeted Tissue kit & NGS** | **Sample Usage** |
| --- | --- | --- | --- | --- |
| NSCLC_004 | Adenocarcinoma | IIA | COSM6224 | Analytical Accuracy |
| NSCLC_006 | Squamous cell carcinoma | IIIB | No hits | Analytical Accuracy |
| NSCLC_007 | Neuroendocrine/large cell neuroendocrine carcinoma | IIIA | COSM516 | Analytical Accuracy |
| NSCLC_010 | Neuroendocrine/large cell neuroendocrine carcinoma | IIB | COSM520 | Analytical Accuracy |
| NSCLC_013 | Adenocarcinoma | Not known | No hits | Analytical Accuracy |
| NSCLC_018 | Adenocarcinoma | IIA | COSM6223 | Analytical Precision |
| NSCLC_021 | Adenocarcinoma | IB | No hits | Analytical Accuracy |
| NSCLC_029 | Adenocarcinoma | III | No hits | Analytical Accuracy |
| NSCLC_048 | Adenocarcinoma | IIB | COSM6223 | Analytical Precision |
| NSCLC_052 | Squamous cell carcinoma | III | COSM6240; COSM6213 | Analytical Accuracy |
| NSCLC_056 | Adenocarcinoma | IIIA | COSM516 | Analytical Accuracy |
| NSCLC_057 | Adenocarcinoma | IIIA | COSM476 | Analytical Accuracy |
| NSCLC_059 | Squamous cell carcinoma | IIIA | COSM516 | Analytical Accuracy |
| NSCLC_061 | Adenocarcinoma | IIIB | COSM6224 | Analytical Accuracy |
| NSCLC_062 | Adenocarcinoma | IIIA | COSM6224 | Analytical Precision |
| NSCLC_063 | Adenocarcinoma | IIIA | COSM6241 | Analytical Accuracy |
| NSCLC_069 | Adenocarcinoma | IIA | EML4/ALK* | Analytical Accuracy |
| NSCLC_079 | Adenocarcinoma | IVB | COSM12376 | Analytical Accuracy |
| NSCLC_081 | Adenocarcinoma | IIB | COSM13428 | Analytical Accuracy |
| NSCLC_084 | Adenocarcinoma | IIIA | COSM6224 | Analytical Accuracy |
| NSCLC_088 | Adenocarcinoma | IA | COSM553 | Analytical Accuracy |
| NSCLC_090 | Squamous cell carcinoma | IB | EML4/ALK | Analytical Accuracy |
| NSCLC_094 | Adenocarcinoma | IB | COSM6223 | Analytical Accuracy |
| NSCLC_098 | Adenocarcinoma | IIIA | COSM476 | Analytical Accuracy |
| NSCLC_103 | Adenocarcinoma | IIIA | COSM6224 | Analytical Accuracy |
| NSCLC_111 | Adenocarcinoma | IA | COSM476 | Analytical Accuracy |
| NSCLC_112 | Adenocarcinoma | IA | COSM6223 | Analytical Accuracy |
| NSCLC_119 | Adenocarcinoma | IIB | COSM516 | Analytical Accuracy |
| NSCLC_122 | Adenocarcinoma | IB | COSM6223 | Analytical Accuracy |
| NSCLC_126 | Adenocarcinoma | IA3 | COSM516 | Analytical Accuracy |
| NSCLC_130 | Large cell carcinoma | IIB | COSM522 | Analytical Accuracy |
| NSCLC_131 | Adenocarcinoma | IIIB | MET exon 14 skipping | Analytical Accuracy |
| NSCLC_134 | Adenocarcinoma | IV | EML4/ALK | Analytical Accuracy |
| NSCLC_141 | Adenocarcinoma | IIIA | EML4/ALK* | Analytical Accuracy |
| NSCLC_150 | Adenocarcinoma | IIIA | COSM6225 | Analytical Accuracy |

**FFPE lung tissue samples from patients with a confirmed NSCLC diagnosis.** Shown are the clinical characteristics associated with each sample, and how each sample was used during the analytical validation. *Fusion breakpoint not predicted to be detected by ASPYRE-Lung.

**S5 Table**

|  | | **Run#** | 1 | 2 | 3 | 4 |  |
| --- | --- | --- | --- | --- | --- | --- | --- |
|  |  | **Instrument** | 1 | 2 | 1 | 2 |  |
|  |  | **Operator** | 1 | 1 | 2 | 2 |  |
|  |  | **Reagent** | 1 | 2 | 2 | 1 |  |
| **Sample ID** | | **Day** | 1 | 2 | 3 | 4 |  |
| **DNA** | **RNA** | **Expected Variant** | **Replicates with expected call / total replicates** | | | | **Total** |
| HBF_0389 | HBF_0414 | Negative | 2/2 | 2/2 | 2/2 | 2/2 | 8/8 |
| HBF_0391 | HBF_0416 | Negative | 2/2 | 2/2 | 2/2 | 2/2 | 8/8 |
| HBF_0392 | HBF_0419 | Negative | 2/2 | 2/2 | 2/2 | 2/2 | 8/8 |
| FFPE_018 | FFPE_018 | COSM6223 | 2/2 | 2/2 | 2/2 | 2/2 | 8/8 |
| FFPE_048 | FFPE_048 | COSM6223 | 2/2 | 2/2 | 2/2 | 2/2 | 8/8 |
| FFPE_062 | FFPE_062 | COSM6224 | 2/2 | 2/2 | 2/2 | 2/2 | 8/8 |

**Expanded Results from testing analytical precision.** Results from testing nine clinical samples across four runs using two different instruments, two different reagents lots, by two different operators over four days.
